# Single molecule characterization of the binding kinetics of a transcription factor and its modulation by DNA sequence and methylation

**DOI:** 10.1101/2021.05.19.444789

**Authors:** Hadeel Khamis, Sergei Rudnizky, Philippa Melamed, Ariel Kaplan

## Abstract

The interaction of transcription factors with their response elements in DNA is emerging as a highly complex process, whose characterization requires measuring the full distribution of binding and dissociation times in a well-controlled assay. Here, we present a single-molecule assay that exploits the thermal fluctuations of a DNA hairpin, to detect the association and dissociation of individual, unlabeled transcription factors. We demonstrate this new approach by following the binding of Egr1 to its consensus motif and the three binding sites found in the promoter of the *Lhb* gene, and find that both association and dissociation are modulated by the 9 bp core motif and the sequences around it. In addition, CpG methylation modulates the dissociation kinetics in a sequence and position-dependent manner, which can both stabilize or destabilize the complex. Together, our findings show how variations in sequence and methylation patterns synergistically extend the spectrum of a protein’s binding properties, and demonstrate how the proposed approach can provide new insights on the function of transcription factors.

## Introduction

Already from the seminal work of Jacob and Monod^1^, a central regulatory role was recognized for the equilibrium occupancy of response elements by TFs, which depends on their concentration, affinity, and, in eukaryotes, the chromatin accessibility of their binding sites^2,3^. However, recent studies have highlighted the importance of transient TF-DNA interactions, and indicate that the *kinetics* of TF binding and dissociation play an important role in regulating transcription *in vivo*^4–11^. In particular, the importance of the TF residence time in determining the transcriptional burst duration^6,10,11^ and size^6,10^ has been demonstrated. It has also been shown that association and dissociation are complex processes^12–19^, that cannot be described as simple second- and first-order reactions, respectively. TFs find their targets by coupling three-dimensional diffusion in the solution with one-dimensional diffusion while bound nonspecifically to DNA^20^, and most are unable to bind nucleosomal DNA, so the association time is affected by the genomic context of the binding site^21,22^ and by a kinetic competition with nucleosomes^23^. On the other side, broad distributions in the residence time of individual TFs were observed in vivo^8^ and also higher-order phenomena, such as dissociation facilitated by other TFs^13,15^ or nucleosomes^24^. Hence, given that TF binding to promoters and enhancers is a central event in the initiation of transcription, whose perturbation is linked to different disease states^25^, quantitative measurements of the binding and dissociation times of TFs are of utmost importance. However, the complexity described above can be obscured by the ensemble averaging limitations of traditional biochemical methods, stressing the importance of following the binding reaction at the single-molecule level to reveal the full distribution of binding and dissociation times. Indeed, much of the evidence for the importance of the binding kinetics, and measurements of kinetic rates, are based on single-molecule tracking experiments, which can follow the motions of nuclear factors in living cells^26,27^. Although very powerful, these experiments also suffer from important limitations^28^: First among them is are the need to label the protein of interest, which can result in a perturbation of its diffusional and binding properties^29,30^. In addition, their typical resolution is much larger than the size of the protein and its binding site. It is also challenging to uncouple the dynamics originating from the chromatin itself and that from the chromatin-binding protein. And, photobleaching and motions out of the focal plane prevent measurements of long trajectories^12^. In vitro single-molecule methods based on fluorescence detection allow for a controlled environment^31^, and have revealed important aspects of the binding of proteins to chromatin^24,32–34^, but also require labeling and are limited to short DNA sequences.

Here we introduce a new and complementary approach to characterize the binding kinetics of unlabeled TFs in a single molecule biophysical assay that can also gradually accommodate the complexity of the chromatin in real genes. Using DNA unzipping with optical tweezers, we exploit the thermal fluctuations of DNA as a sensor to follow, in real-time, the binding and dissociation of individual TFs to a binding site in its native sequence environment. We demonstrate our approach with Egr1, an inducible transcription factor responsible for regulating a variety of genes^35^, and the *Lhb* gene, which encodes for the Luteinizing Hormone beta subunit and is expressed in gonadotrope cells^36^. The DNA binding domain of Egr1 contains 3 zinc fingers (ZFs), each interacting with 3 DNA base pairs, making a 9 bp binding motif^37^. Genome-wide studies obtained a consensus motif (CGCCCACGC) to which Egr1 binds with high affinity^38^. However, most functional Egr1 binding sites in the genome deviate from this consensus, and these differences are gene-specific and evolutionarily conserved. This is also the case for *Lhb, which* contains 3 different binding sites in its proximal promoter. In a previous work^39^, we showed that their specific sequence, and the conserved sequences flanking the binding sites, modulate the binding affinity and the conformation of the bound protein, suggesting that a functionally important and sequence-specific spectrum of conformations exist for Egr1. Here, we study how the sequence of the different sites modulates the binding kinetics. We found that the DNA sequence, at the core binding site and flanking it, modulates both binding and dissociation rates. Next, given that hypermethylation of Egr1 binding sites is associated with developmental defects^40–43^ and that the proximal promoter of *Lhb* is hypomethylated in gonadotrope cells^44^, we monitored how methylation of the *Lhb* sites, each with a distinct CpG methylation pattern, affects the binding kinetics. Our data show that methylation of CpGs modulates only the protein’s residence time on the DNA. This modulation depends on the sequence of the DNA and the position of the methylation site and can both stabilize and destabilize the complex. Together, our results demonstrate the strength of the proposed method and highlight the versatility of DNA sequence and CpG methylation as a mechanism for TF binding regulation.

## Results

### Real-time measurements of the binding and dissociation of individual TFs

To follow the binding and dissociation of a single, unlabeled Egr1 protein to its binding site, we use optical tweezers to monitor the spontaneous fluctuations of a partially unzipped DNA molecule. Each of the strands of a ∼400 bp DNA “sample” containing the consensus binding site for Egr1, is attached to a long (∼2000 bp) dsDNA “handle” (Figure 1A). The other end of each handle is bound to a polystyrene bead held in one of the traps of a dual trap optical tweezers setup (Figure 1A). By steering one of the traps away from the other, creating a tension above 17-18 pN, the DNA sample is unzipped in a sequence-dependent pattern, consisting of successive drops in force that are concomitant with increases in the overall extension (Figure S1A), which represent the cooperative opening of a few base pairs (Figure S1B). When the unzipping fork reaches the vicinity of the TF binding site, we stop the movement of the traps and hold them at a constant distance from each other. The DNA sample, which is under tension, exhibits spontaneous “breathing” dynamics^45^, i.e., the thermally driven opening and closing of a short (25-30 bp) DNA segment (Figure 1B). In general, due to the differences in binding energy between CG and AT pairs, these fluctuations populate only two states, which we term “open” and “closed”. Next, exploiting a laminar flow cell, the fluctuating DNA molecule is moved to a different region where it is exposed to a solution containing Egr1 (10 nM, unless specified otherwise). Binding of the protein to its binding site stabilizes the closed state, momentarily suppressing the breathing, which reappears upon dissociation of the protein (Figure 1A,C). Since the typical time the oscillations remain suppressed is much longer than the time spent in the closed state by a protein-free DNA molecule (Figure S1C-D), we are able to set a threshold (*t*_*TH*_= 150 ms) and reliably identify the protein-bound state as a stable closed state whose duration is longer than *t*_*TH*_. No bound events are identified in the absence of proteins (Figure 1B, Figure S1C) or for DNA sequences that do not include a binding site for Egr1. Identification of multiple bound and free states allows us to extract the mean binding and residence time, and their reciprocals, the observed binding and dissociation rates, 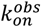 and 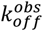.

**Figure 1.**
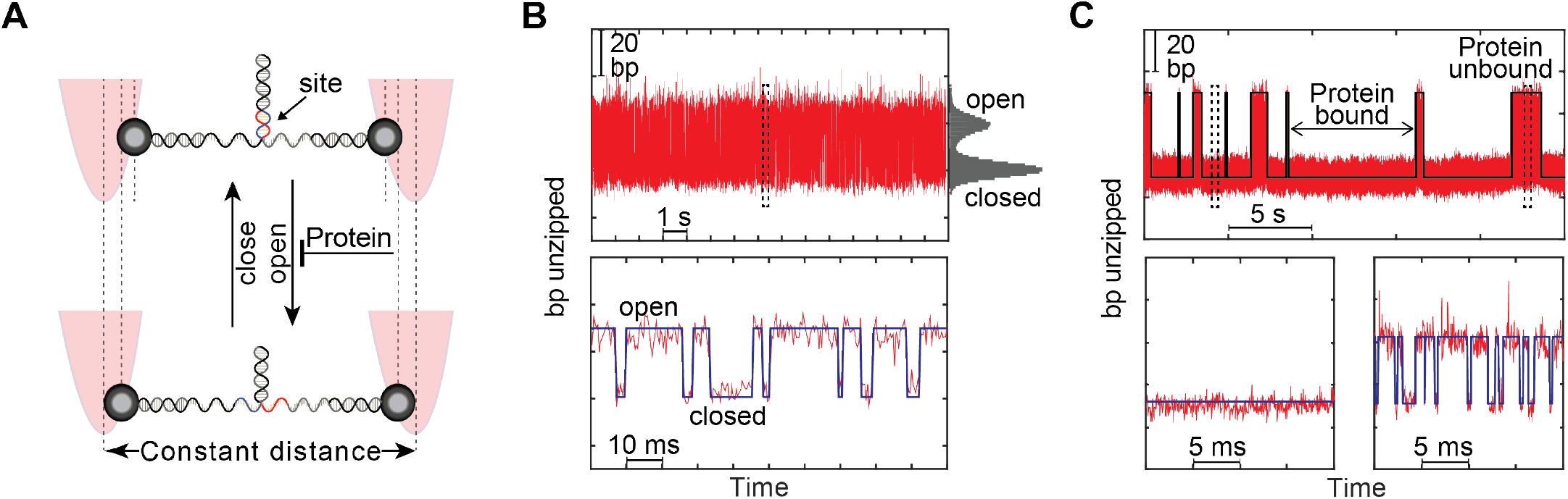
Real-time measurements of residence and binding times. **(A)** Schematic description of the experiment. The DNA construct is attached from both sides to polystyrene beads trapped in optical traps. The distance between the traps is gradually increased, unzipping the DNA until reaching the vicinity of a binding site for Egr1, and then kept at a constant value. A segment of ∼25-30 bp in the fork, which includes the binding site, thermally fluctuates between an open and a closed state. **(B)** Breathing fluctuations in the number of base pairs unzipped. The histogram on the right side corresponds to the probability of populating the two states, open and closed. The lower panel shows a close up look at the dashed region in the upper one. **(C)** Breathing fluctuations in the presence of Egr1. Binding of the protein suppresses the DNA fluctuations and is detected as a long and stable closed state, while the fluctuating regions correspond to unbound protein states. The lower panel provides a closer look into the bound (left dashed box in the upper panel) and unbound (right dashed box) protein states, respectively. Data in (B-C) is shown unfiltered, as sampled at 2.5kHz.

### Egr-1 binds passively to dsDNA, and exhibits two dissociation regimes

Notably, the ability to control the force applied on the DNA opens also the possibility to shed light on the *mechanism* of binding. In principle, Egr1 binding can occur through one of three pathways (Figure 2A): First, it can bind passively to the already closed DNA. Next, it can bind to the open ssDNA and catalyze closing of the duplex. Finally, the protein may bind to open ssDNA and wait for the DNA to close spontaneously. Assuming that the DNA thermal fluctuations are in rapid equilibrium relative to Egr1 binding, each one of these pathways predicts a different functional dependence on the probability *p*_*closed*_ of the DNA being close (Supplementary Discussion). The first pathway suggests that 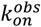 will increase linearly, while the other two suggest 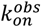 will decrease linearly with *p*_*closed*_. In order to elucidate which one is the dominant binding pathway, we exploit the fact that changing the distance between the traps (Figure 2B), and hence the applied tension on the fork, results in a very sensitive modulation of *p*_*closed*_, from ∼1 to ∼0 in a ∼1 pN range (Figure 2C-F). These measurements reveal a linear dependence of 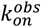 on *p*_*closed*_ (Figure 2G), which is an indication that Egr1 binds only to the closed DNA, exploiting a momentary and thermally driven closing of the fork (Figure 2A). Interestingly, the inhibition of binding by thermal fluctuations, together with the suppression of these fluctuations by a bound protein, offer an interesting mechanism for binding cooperativity between different TFs.

**Figure 2.**
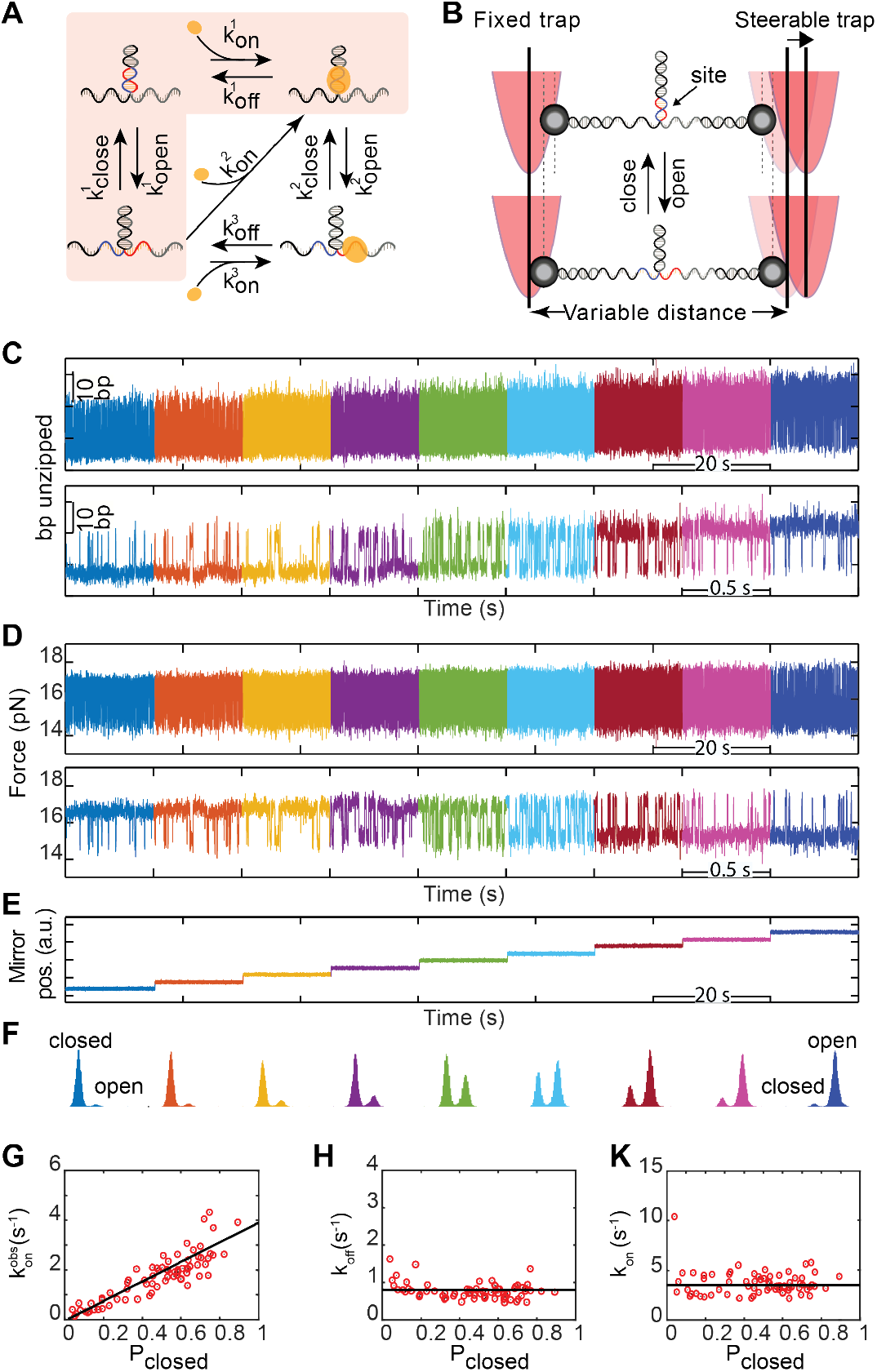
Egr1 binds passively to DNA. **A)** Possible association paths of Egr1. **(B)** Illustration of the experiment. The position of the steerable trap is gradually shifted in a controlled manner to adjust the tension on the tether. **(C-E)** Number of base pairs unzipped (C) a force (D) obtained from the DNA breathing. Each color corresponds to a different distance between the two traps (E) characterized by a different probability to occupy the closed state. The lower panels in (C,D) are a zoom-in to 0.5 s at each condition from the same experiment. Data are shown as sampled, at 2.5 kHz. **(F)** Histograms of the extension for each condition. The lower value in the histogram corresponds to the closed state and the higher value to the open state. **(G)** Observed association rates as a function of *p*_*closed*_. **(H)** Observed dissociation rate as a function of *p*_*closed*_.**(K)** Intrinsic association rate, obtained by dividing the observed association rate by *p*_*closed*_, as a function of *p*_*closed*_. Data in (E-G) correspond to site -1 in the *lhb* proximal promoter. N=11 DNA molecules, 1818 and 1790 detected times, for binding and residence, respectively

In contrast to the association rate, the observed dissociation rate is independent of the tension on the fork (Figure 2H). However, since our method is based on monitoring the DNA fluctuations, and thus requires 0 < *p*_*closed*_< 1, the range of forces we can probe is very narrow (Figure 2D). To widen this force range and elucidate whether the residence times we measure are affected by the applied tension, we develop a new assay (Figure 3A). First, we detect Egr1 binding via the suppression of the breathing fluctuations. Once a binding event is detected, we rapidly (with a 150±50 ms response-time) move one of the traps in a single step to increase the applied tension. The distance between the two traps is then held constant until the force drops, indicating protein dissociation and revealing the bound state’s lifetime at the specific force probed. Finally, the trap is restored to its original position allowing for additional measurement rounds. Using this approach, we can explore the residence time at an extended force range (Figure 3B,C). Theoretical models for such dynamic force spectroscopy experiments^46,47^ predict an exponential decrease in the dissociation kinetics as a function of the force, allowing the extraction of the transition state’s location and energy. However, our data do not show a simple exponential decay, but rather a turnover from a force-independent range to a force-dependent one that fits well to an exponential function (Figure 3C). This behavior seems to be independent of the sequence of the binding site (Figure 3D). This type of turnover has been observed in studies probing protein unfolding under force^48^ and suggests the existence of two pathways for dissociation: a force-independent one and a forced-induced dissociation pathway with a different rate. Notably, given that unzipping the DNA partially mimics the progression of polymerases and helicases on the DNA, it is possible that these different dissociation pathways play a role in ensuring stable binding of the protein to DNA in the absence of a translocating enzyme, but efficient and rapid dissociation in their presence.

**Figure 3.**
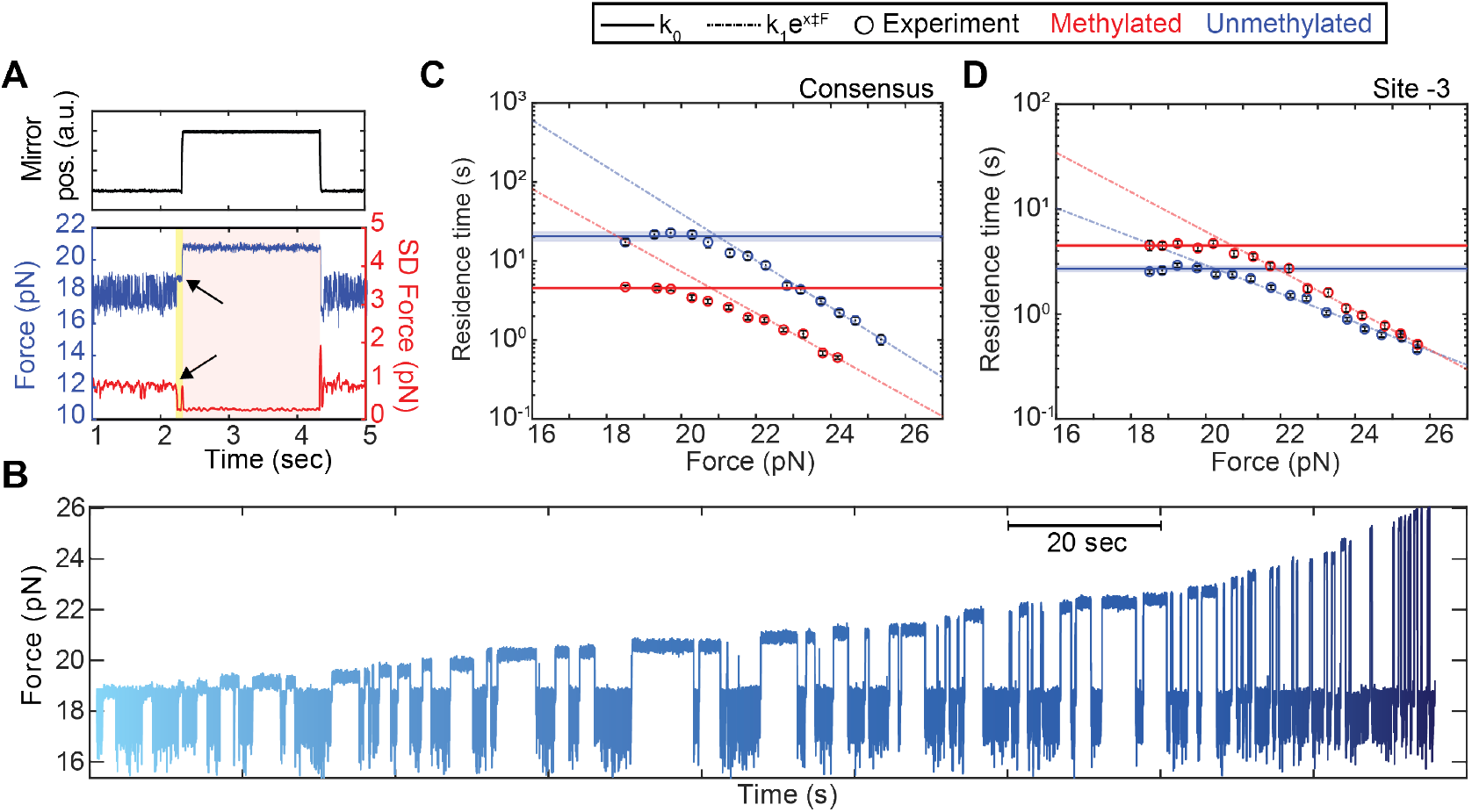
Two dissociation pathways exist for Egr1. **(A)** Illustration of the dissociation under force assay. Top panel: Mirror position vs. time. Bottom panel: external force (left Y-axis, blue), and standard deviation of the force (right Y-axis, red) vs. time. The experiments response time is shaded in yellow. Times at which Egr1 is bound are shaded in red. Data were filtered to 200 Hz (Butterworth). **(B)** Representative binding events at various applied forces (different shades of blue). Data shown as sampled, at 2.5 kHz. **(C-D)** Residence time vs. applied force for unmethylated (blue) and methylated (red) DNA for the consensus site (C) and site -3 in *Lhb* (D). The average of data points at forces below 20 pN is shown as a horizontal line ± standard deviation. An exponential decay fit for the data higher than 20 pN is shown in dashed lines.

Since the turnover takes place at forces that are higher than those used in the fluctuation’s measurements of Figure 1, these experiments reveal the inherent, or force-free association kinetics. With the typical concentrations used in this study, hundreds of binding and dissociation events can be identified in a single trace and used to calculate the cumulative distribution function (CDFs) for the bound state duration (Figure S1E). In addition, we calculate the CDF for the inherent association time, obtained by dividing the measured times by the instantaneous *p*_*closed*_ (Figure 2K, Figure S1F). While, in general, the shape of these distributions may reveal the existence of several underlying processes controlling association and dissociation, in this case, we find that both are well-fitted by exponential functions, which allows us to extract the inherent, force-free binding and dissociation rates, *k*_*on*_ and *k*_*off*_, respectively. Notably, although the measured *k*_*off*_ is highly consistent between different experimental sessions (Figure S2A), we do observe some variability in the measured *k*_*on*_, likely as a result of small changes over time in the effective concentration (Figure S2B). To minimize this effect, we compare in what follows only values of *k*_*on*_ that were obtained in the same experimental session. With this approach, we can now ask how these rates are modulated.

### The sequence and context of the binding site modulate the binding kinetics

In a previous work, using DNA unzipping to disrupt Egr1-DNA complexes at equilibrium, we reported significant differences in affinity between the consensus site and the three evolutionary-conserved binding sites at the proximal promoter of the *Lhb* gene. With the approach described here, we are now able to characterize the more informative *kinetics* of binding. In other words, while the previous study indicates that *k*_*on*_/*k*_*off*_ is affected by the DNA sequence, we can now probe the effect that the sequence has on *k*_*on*_ and *k*_*off*_ separately. Measurements of the *Lhb* sites in their native genomic context, and the consensus site inserted in the 601 nucleosome positioning sequence^49^ (Figure 4A) reveal that both the dissociation and association kinetic rates are modulated by the identity of the binding site (Figure 4B-D). To uncouple between the potential effect of the core binding sequence and that of the sequences surrounding the binding sites, we also probe the *Lhb* sites when inserted in the 601 context and the proximal promoter of the *Cga* gene^50^ (Figure S3A). Differences in *k*_*off*_, which are also affected by the flanking sequences (Figure S3B), are consistent with a modulation in the structure of the complex^39^. Differences in *k*_*on*_ (Figure S3C), suggest that binding is not diffusion-limited but includes an additional step of transition into a tightly bound complex, and that this step occurs at a different rate for each site. Interestingly, this “isomerization” step is also affected by the sequences flanking the binding site, but differences between the sites are present even when inserted in a common context (Figure S3C), indicating that both the core and flanking sequences can affect isomerization. A simple two-state model (Figure S4) allows estimating their relative contributions (Tables S1-5), and reveals a non-additive effect for mutations in the core binding site, in contrast to previous studies^51^ (Supplementary Discussion). Overall, our data stress the importance of the genomic environment as a modulator of binding kinetics.

**Figure 4.**
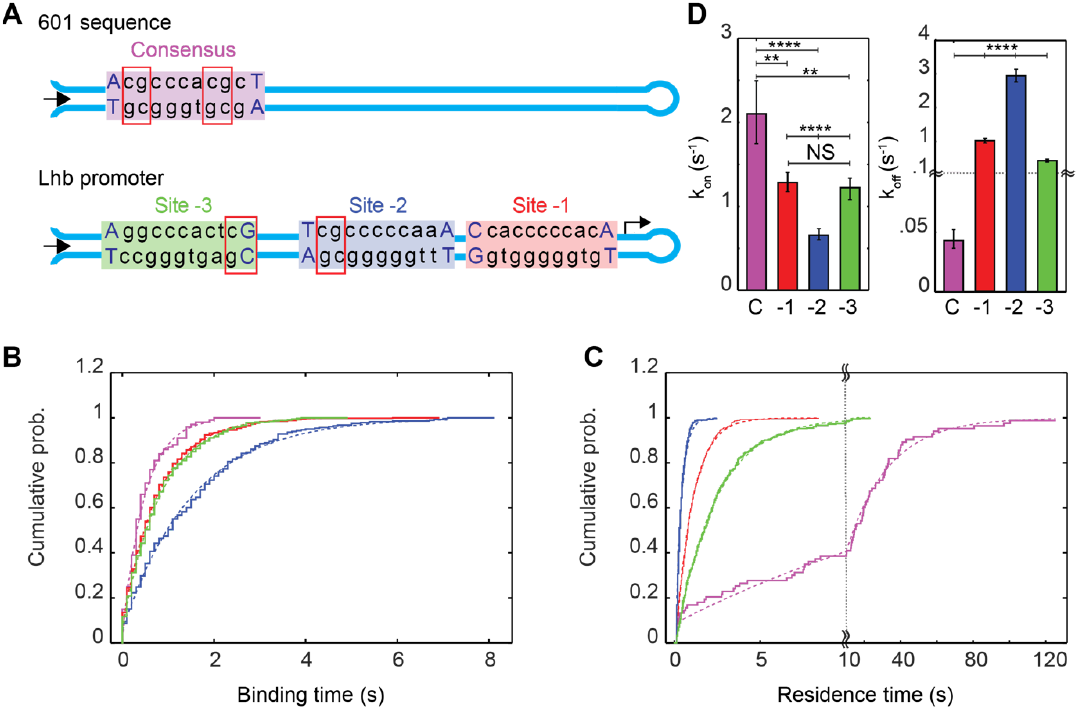
The sequence of Egr1 binding site modulates both its association and dissociation kinetics. **(A)** DNA constructs containing the consensus binding site inserted in the 601 DNA sequence (top) and the *Lhb* proximal promoter with its 3 identified sites (bottom). CpG sites are highlighted with red squares. The unzipping direction is indicated as a black arrow. **(B-C)** Cumulative probability distribution functions for the binding (B) and the residence (C) times, for the consensus (magenta), site -3 (green), site -2 (blue) and site -3 (red) at [Egr1] = 10 nM. A bin size of 0.2 s was used for the binding time. For the residence time, a variable bin size was used: 2 s for the consensus, 0.5 s for site -3 and -1, and 0.1 s for site -2. The data presented were obtained in a single session. N=6 molecules, 100 and 83 binding and dissociation events, respectively, for the consensus; 5, 511 and 508 for site -1; 4, 278, 296 for site -2; and 4, 359, 353 for site -3. A single-exponential fit is presented as a dashed line. **(D)** Association (left) and dissociation (right) rates. Data are shown as the average of the rates extracted from the fit ± the mean of the errors, taken from 6 different sessions conducted at the same Egr1 concentration (10 nM). \*\*\*\**P* < 0.0001, ***P < 0.001, *P < 0.05, Student’s t-test.

### Methylation induces a sequence- and position-dependent modulation of the binding kinetics

The conserved differences in the core sequences of the *Lhb* binding sites also imply that they harbor different potential CpG methylation sites, which can add an additional, and differential, layer of regulation to Egr1 binding to the different sites. Hence, we ask whether methylation can also modulate the binding kinetics, and if so, which of the kinetic rates is affected. We first probe a construct containing the consensus motif (Material and methods, Figure S5), which contains two methylation sites, one on each side of the motif (Figure 4A), and characterize the distributions of binding and residence times (Figure 5A,B). Interestingly, while methylation by M.SssI methyltransferase does not affect *k*_*on*_ (Figure 5A), we observe a remarkable sensitivity of *k*_*off*_ for the methylation state (Figure 5B), which represents a 5-fold decrease in transcription factor residence time, and a reduction in the binding energy of ∼ 1 kCal/mol (Supplementary Discussion, Tables S6). This is in stark contrast with previous works^52,53^, although likely the result of using a long DNA segment in our study (Supplementary Discussion, Figure S6). Interestingly, an irreversible unzipping experiment, where we continuously increase the force to induce dissociation of a bound protein^39^ reveals no differences in the binding position upon methylation, but significant differences in the breaking force (Figure S7), which suggests a different conformation of the protein when bound to methylated DNA.

**Figure 5.**
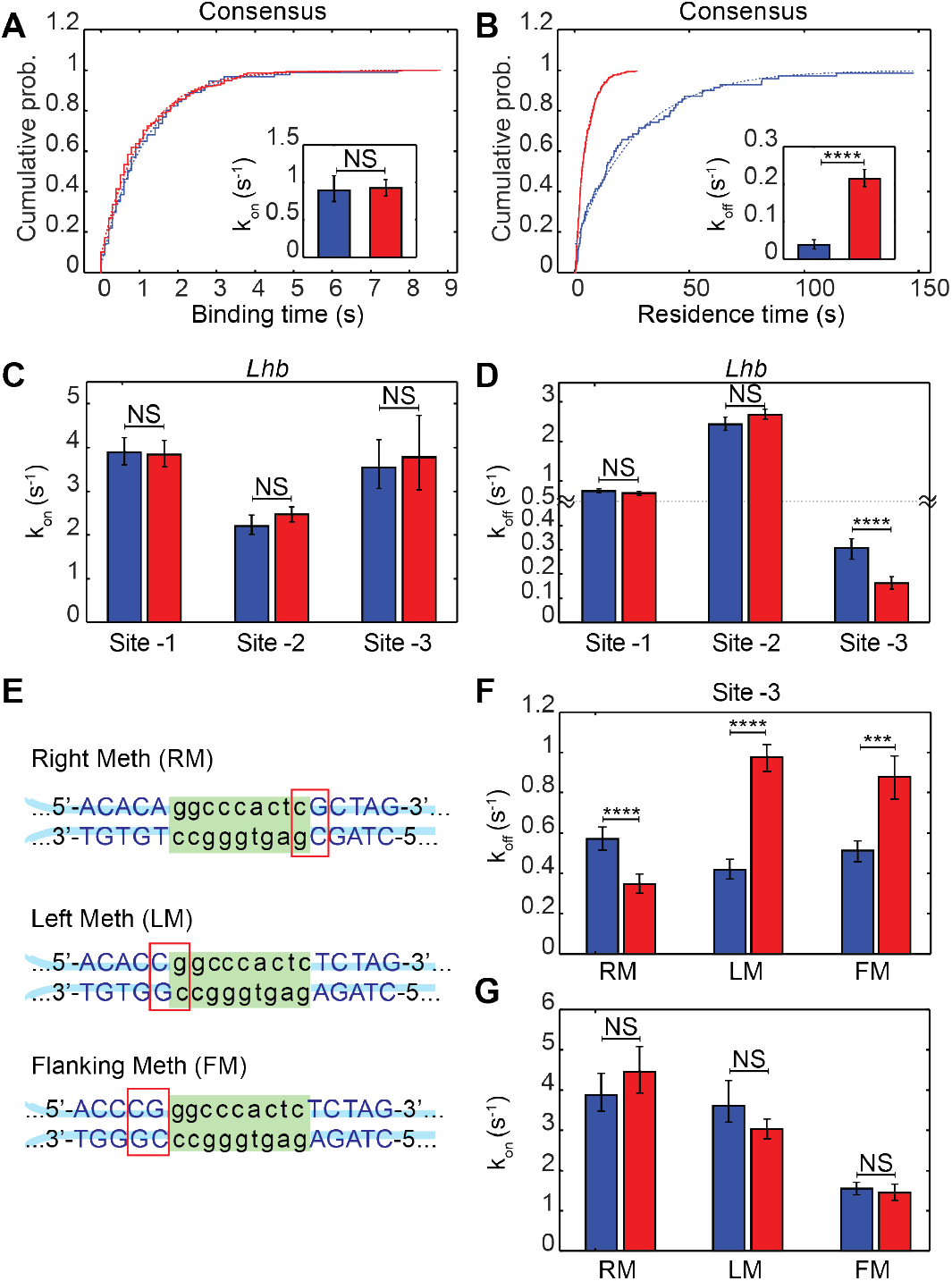
Methylation modulates the residence time in a sequence- and position-depending manner. **(A-B)** Cumulative distributions of the binding (A) and residence (B) times for unmethylated (blue) or methylated (red) consensus binding site. The dashed line shows a single exponential fit. The inserts show the rate obtained from the fit ± 95% confidence interval obtained by bootstrapping (resampling the data 100 times, with replacement). The experiment is done at 1 nM of Egr1. N = 6 molecules, 91 binding events, and 70 dissociation events for the unmethylated case; 4, 304, 299 for the methylated one. **(C-D)** Association (C) and dissociation (D) rates for the unmethylated (blue) and methylated (red) 3 sites in the *Lhb* promoter, calculated and presented as in A and B. The experiment is done at 10 nM Egr1. N = 3 molecules, 544 and 535 detected binding and dissociation events, respectively, for site -1 unmethylated; 4, 759, 744 for site -1 methylated; 3, 449, 460 for site -2 unmethylated; 5, 707, 716 for site -2 methylated; 3, 210, 197 for site -3 unmethylated;5, 124, 112 for site -3 methylated. **(E)** Sequence for site -3 from the *Lhb* promoter, integrated into a new context with a CpG inside the binding site, in its left (LM) and right (RM) sides, or a CpG outside the 9bps core of the binding site (FM). **(F-G)** Dissociation (F) and binding (G) rates for the constructs on (E), calculated and presented as in A and B. The experiment is done at 10 nM Egr1 for constructs RM & LM, and at 20 nM for FM. N = 4 molecules, 263 and 264 detected binding and dissociation events, respectively, for unmethylated RM; 3, 202, 196 for methylated RM; 3, 276, 269 for unmethylated LM; 4, 489, 490 for methylated LM; 5, 302, 302 for unmethylated FM; 3, 173, 178 for methylated FM. ****P < 0.0001, ***P < 0.001, NS P>0.05, Kolmogorov-Smirnov’s t-test. Each methylated (red) and unmethylated (blue) pair was collected at the same Egr1 concentration and at the same session for accurate comparison.

Next, when we repeat our assay using the *Lhb* sites, no significant effect is observed upon treatment with M.SssI for the binding time of any of the sites, consistently with the results for the consensus site (Figure 5C). The residence times for site -1, which has no CpG sites in its core binding site or near flanking sequences, and site -2, which has a single methylation site, are also insensitive to the methylation reaction (Figure 5D). However, site -3, which also contains a single CpG site but on an opposite orientation to that of site -2 (Figure 4A), shows a two-fold increase in residence time, indicating that methylation makes the complex more stable, by -0.4 ± 0.2 kCal/mol (Figure 5D, Table S6). Interestingly, the modulation provided by the methyl group in this case is opposite to that of the consensus motif, indicating that methylation of Egr1 binding sites can have both stabilizing and destabilizing effects.

To clarify whether the differences observed for site -3 and the consensus (stabilization and destabilization, respectively) are the result of their different core sequence, or their different flanking context, we measure the kinetics of site -3 inserted into the same context as the consensus motif (Figure 4E-G, construct RM). Consistent with the previous result, a significant decrease in *k*_*off*_ and no effect on *k*_*on*_ are observed upon methylation (Figure 5F,G), suggesting that the core composition, but not the flanking context is responsible for the observed effect of DNA methylation. Since Egr1 binds asymmetrically to its binding motif, and demonstrates a greater sensitivity to a perturbation from the side of ZF3^39^, we hypothesize that the position of the methyl group relative to the ZFs may dictate the stabilizing or destabilizing effect of the methylation. Hence, we eliminate the CpG adjacent to ZF3 in site -3 and insert a new one adjacent to ZF1, without changing the 9 bp core sequence (Figure 5E, construct LM). Remarkably, the methylation effect on the stability is now reversed, resulting in destabilization of the complex (Figure 5F) and providing support for our hypothesis. Finally, shifting the methylation site on the LM construct to be outside the core binding site, but immediately adjacent to ZF1 (Figure 5E, construct FM), results in a similar destabilization effect upon methylation (Figure 4F). Hence, the same binding site can exhibit different and sometimes opposite responses to methylation, depending on the position of the CpG site, both in the core and flanking sequences. Together, the large but static modulation range provided by differences in core and flanking sequence, and the dynamic control of dissociation rate by CpG methylation, likely contribute to the broad functionality of a single TF.

## Discussion

Here we demonstrate a single molecule biophysical assay that allows to follow, in real-time, the binding and dissociation of individual, unlabeled TFs to DNA. Our method is based on monitoring the thermal fluctuations of DNA, which are suppressed upon protein binding. With the ability to obtain the full probability distribution for the binding and residence times of a TF to a known site, while at the same time offering the possibility to gradually increase the complexity of the system and follow binding in the vicinity of nucleosomes^54^ and chromatosomes^55^, we expect this method to offer important mechanistic insights to complement the information obtained from live cell measurements.

We demonstrate our approach by studying binding of Egr1 to the *Lhb* proximal promoter. In line with our previous work, which showed that the equilibrium properties of binding are modulated by the conserved deviations found in the sequence of the *Lhb* sites and the conserved sequences around them, here we show that both the kinetic rates, *k*_*on*_ are *k*_*off*_, are separately modulated by these factors. Different binding sites within the same DNA segment, measured at the same Egr1 concentration, exhibit differences in *k*_*on*_, which characterizes the typical duration needed for Egr1 to bind in a stable conformation to its site and inhibit its fluctuations. On the other hand, *k*_*on*_ depends also on the concentration (Figure S8), suggesting that the binding process has more than one step, with only the first one involving diffusion of the TF to the vicinity of the binding site. Previous studies showed that Egr1 is able to perform 1D diffusion on the DNA^56,57^, thus increasing the speed at which it finds its binding site by “facilitated diffusion”^20^. It does so by exploiting two different modes of DNA binding: the “search” mode, with two ZFs attached to the DNA, has low binding energy and allow fast and efficient DNA scanning, while the “recognition mode”, where all three ZFs are bound to the DNA, has higher binding energy^56^. The existence of these two conformations enables Egr1 to overcome the “speed-stability” paradox, i.e., the contradicting needs for both efficient sliding on non-specific DNA and stable binding to a specific site^58^. Since our experiments only detect stably bound proteins, one possible interpretation for the context-dependence of *k*_*on*_ is that the observed isomerization step involves, at least partially, 1D diffusion into the binding site. However, differences in *k*_*on*_ were observed also for different sites inserted in a common context, indicating that this effect may only partially explain the results. While it is also possible that a *transition* between the search mode and the recognition mode is modulated by the sequence of the site itself, a thorough elucidation of the nature of the different steps involved in reaching the binding site and stably binding to it will require additional studies. Remarkably, differences in residence time as large as 40-fold were observed as a result of 3 bp differences in the binding site, which is consistent with studies showing a broad, power-law distribution in the residence time for other transcription factors in vivo. Moreover, the high degree of sequence conservation observed for the binding sites of Egr1 suggests an important role played by these differences in residence time in the differential regulation of the many genes controlled by Egr1.

Different sequences correspond to different potential methylation patterns, and while CpG methylation does not affect the kinetics of Egr-1 binding, our data shows that it modulates the dissociation of the protein. This effect is sensitive to the position of the CpG in the binding site or its vicinity and can result in both stabilization and destabilization of the complex. Interestingly, a structural study of CTCF, which contains an array of 11 ZFs, also revealed sensitivity for CpG methylation in a position-dependent manner^59^. How does a methyl group in the binding site affect the stability of the complex? While one possible mechanism is steric hindrance by the methyl group, the effect observed for a methylated CpG outside the core binding site, and the fact that methylation can stabilize as well as destabilize the bound complex, suggest that part of the effect is related to changes in the local structure of DNA. Although similar crystal structures were reported for methylated and unmethylated DNA^60^, methylation was shown to affect the flexibility^61,62^, hydration structure^63^, and sugar pucker conformations^64^ of DNA. Molecular dynamics simulations showed that methylation increases, locally, the propensity of DNA toward different values of roll and propeller twist, and that the position of the modification and the local sequence context has significant effects on the amount of structural variation observed^65^. The formation of protein-DNA complexes involves changes, or distortions, in the structure of DNA, and these are clearly accompanied by an energetic cost. A recent study highlighted this by showing that, in many cases, the affinity of TFs for DNA that includes mismatches, and is therefore distorted *a priori*, is higher than that for non-perturbed DNA^66^. This increase in affinity correlates with distortions that resemble those that are observed in the DNA when it is part of a bound complex. A similar mechanism of “pre-paying” the energetic cost of binding by distorting the DNA, may explain the observed effect of the methylation on the binding of Egr1. In addition, DNA molecules are polymorphic and dynamic, locally exploring different conformations, that are also sequence-dependent^67–69^, and methylation may affect the dynamic equilibrium between different structural states^70–73^, so it is possible that increasing the relative occupancy of the properly distorted state in the ensemble, or modifying the rates of interconversion between different structures, contribute to our results.

Interestingly, we observe a high degree of destabilization upon methylation for the consensus sequence. A very long residence time may be necessary to fulfill a specific function, e.g., functioning as a pioneer transcription factor^32^ or driving cell differentiation^74^. However, such a long residence time may also be toxic for the cell in other circumstances. Hence, methylation of the consensus site may serve as an important switch to prevent this toxicity, while allowing long binding “on-demand”.

In summary, our study demonstrates a novel single-molecule approach that allows characterizing the different factors that affect the binding and dissociation kinetics of a TF, or other DNA binding proteins. With Egr1 as a model, we demonstrate that the sequence of DNA and its methylation state are powerful and versatile modulators of these kinetics. In a broader context, previous studies have shown that significant genetic variants are located in non-coding regions, and that complex phenotypes are the result of altered binding of TFs^75^. Together with previous results showing the importance of the binding kinetics in tailoring the transcriptional response, our results demonstrating that these kinetics are sensitive to small differences in core sequence, environment, and methylation state, may help explain the mechanism by which these phenotypes arise.

## Materialsand Methods

### Reagents

The original plasmid for the DNA binding domain of Egr1, from Dr. Scot Wolfe, was kindly provided by Dr. Amit Meller. The protein was expressed and purified as previously described^39^.

### Optical Tweezers

Experiments were performed in a custom-made double-trap optical tweezers apparatus, as previously described^39,50,76^. Briefly, the beam from an 852 nm laser (TA PRO, Toptica) was coupled into a polarization-maintaining single-mode optical fiber. The collimated beam out of the fiber, with a waist of w_0_=4mm, was split by a polarizing beam splitter (PBS) into two orthogonal polarizations, each directed into a mirror and combined again with a second PBS. One of the mirrors is mounted on a nanometer scale mirror mount (Nano-MTA, Mad City Labs). A X2 telescope expands the beam, and also images the plane of the mirrors into the back focal plane of the focusing microscope objective (Nikon, Plan Apo VC 60X, NA/1.2). Two optical traps are formed at the objective’s focal plane, each by a different polarization, and with a typical stiffness of 0.3-0.5 pN/nm. A second, identical objective collects the light, the two polarizations separated by a PBS, and imaged onto two Position Sensitive Detectors (First Sensor). The beads’ position relative to the center of the trap is determined by back focal plane interferometry^77^. Calibration of the setup was done by analyzing the thermal fluctuations of the trapped beads^78^, which were sampled at 100kHz. Experiments were conducted in a laminar flow chamber (Lumicks), which was passivated following a published protocol^79^ with some modifications. Briefly, we wash the chamber twice by flushing alternately 1M NaOH and Liquinox 1% for 10 min each. Casein (1%) is sonicated and filtered, diluted to 0.2%, flushed into the chamber and incubated in it for 40 min. After the incubation, the system is washed using the working buffer until all the free Casein is flushed. Using this system, we were able to work with very low Egr1 concentrations, ∼ 1nM.

### Molecular construct for single-molecule experiments

The constructs for single-molecule experiments were generated as described previously^50^, with a number of changes. Briefly, TF binding segments containing the *Lhb* promoter sequence were amplified by PCR from mouse genomic DNA, and segments for the non-native contexts (the 601 nucleosome positioning sequence and the *Cga* gene promoter) were amplified from a plasmid (a gift from Daniela Rhodes (MRC, Cambridge, UK)) and mouse genomic DNA, respectively ^49^. Primers used for the amplification reactions are listed in Tables S7. The constructs were digested using DraIII-HF (New England Biolabs) overnight according to the manufacturer’s instructions. A 10 bp hairpin (Sigma) was ligated to the construct using T4 DNA ligase (New England Biolabs), in a reaction with 1:10 molar excess of the hairpin, at 16 °C. The construct was subsequently digested overnight with BglI (New England Biolabs). Methylation of the constructs was done using CpG Methyltransferase (M.SssI) and s-adenosylmethionine (sam) from New England Biolabs. The methylation efficiency was tested in a restriction reaction with FauI (New England Biolabs), whose activity is abolished for methylated DNA (Figure S5).

We generated two ∼2000-bp DNA handles, each incorporating a specific tag (double digoxygenin and biotin), using commercially purchased 5’ modified primers (Table S8) in a standard PCR reaction, using bacteriophage lambda DNA as the template. The other two primers(Table S8) were designed to contain repeats of three DNA sequences recognized by single-strand nicking enzymes: Nt.BbvCI and Nb.BbvCI (both from New England Biolabs) on the biotin-tagged handle and the digoxygenin-tagged handle, respectively. The nicking enzymes generated 29 nt complementary overhangs on each handle. Handles were mixed at equal molar ratios for DNA annealing, creating a ∼4000 bp fragment of annealed DNA handles. A ∼350 bp dsDNA alignment segment with the sequence of the 601 DNA was prepared using commercially purchased primers (Table S8) in a standard PCR reaction, ligated to the handles, and gel-purified (QIAquick 28706, Qiagen). Binding segments were ligated to DNA handles using a rapid ligase system (Promega) in a 3:1 molar ratio, 30 min at room temperature. The full construct (i.e., handles + alignment segment + TF binding segment) was incubated for 15 min on ice with 0.8 µm polystyrene beads (Spherotech), coated with anti-Digoxigenin (anti-DIG). The reaction was then diluted 1000-fold in binding buffer (10 mM Tris·Cl pH 7.4, 1 mM EDTA, 150 mM NaCl, 1.5 mM MgCl_2_, 1 mM DTT, 3% v/v glycerol and 0.01% BSA). Tether formation was performed *in situ* (inside the optical tweezers’ experimental chamber) by trapping a DNA-bound anti-DIG bead in one trap, trapping a 0.9 µm streptavidin-coated polystyrene beads in the second trap, and bringing the two beads into close proximity to allow binding of the biotin tag in the DNA to the streptavidin in the bead.

### Data analysis

#### Data collection

Data were sampled and stored at 2500Hz. All further processing and analysis were done using Matlab functions. dsDNA was modeled with an Extensible Worm Like Chain model (eWLC), with persistence length 45 nm, inter-phosphate distance 0.34 nm, and stretching modulus 1200pN. For ssDNA we used a WLC model with persistence length 0.67 nm and inter-phosphate distance 0.78 nm. The number of bps unzipped was calculated from the extension measurements by first subtracting the extension of the handles and then dividing by twice the extension of a single ssDNA base at the given tension.

#### Full unzipping experiments

Detection of bound proteins in full unzipping experiments (Figure S1A, blue & Figure S7A) was based on an increase in the force of more than 0.5 pN relative to the median force at the same position for experiments in the absence of Egr1. Applying the same criteria for the data obtained without Egr1 resulted in no binding events detected. Events were classified as belonging to a specific site if the breaking event was located within a 20 bp window relative to the binding site’s expected center. The significance of differences in the breaking forces was assessed with a two-tail Student’s t-test.

#### Measurements of the dissociation and association times

DNA was unzipped until the position of a known Egr1 binding site, and the applied force was further adjusted to result in a given probability of ‘closed’ vs. ‘open’ configuration. The fluctuations in extension were measured for 1 min, after which the construct was moved to a region in the laminar flow chamber containing Egr1. Detection of closed and open states, and hence the time durations for each state, were done using a hidden Markov model, implementing the HAMMY algorithm usually applied to smFRET data^80^. The protein’s binding was identified by a sudden transition to a ‘closed’ state that lasted more than 150 ms. No binding events were detected in the absence of protein (Figure S1C-D), or in the absence of a binding site. The residence time was defined as the time lapsed until the reappearance of the fluctuations.

Using different channels of the laminar flow cell, we can measure the binding kinetics in the same solution for up to 3 different DNA constructs in a single experiment. This allows us to minimize any potential error in the effective concentration of active proteins (Figure S2), and we only compare data taken from the same session. For each session, we build cumulative distribution curves for the residence and binding times, fit them to exponential functions, and extract the association and dissociation rates. The rates are presented as the value obtained from the fit ±95% confidence intervals, obtained from bootstrap by resampling the data 100 times, with replacement.

### Kinetics under force

The DNA was unzipped until reaching the position of Egr1 binding site, and the distance between the traps was then fixed to observe extension fluctuations between the open and the closed states. A drop in the standard deviation of the force, calculated in real-time in 100 ms windows, indicates a protein binding event. Upon detection of the binding event, the voltage in the mirror is instantaneously increased, moving quickly (150±50 ms response-time) the steerable trap to increase the applied force to a predetermined value. The distance between the two traps is then held constant until the force drops, which indicates dissociation of the protein and reveals the bound state’s lifetime at the specific force probed. Next, the mirror voltage, and hence the trap position, are restored to their original values to allow a successive round. Using this protocol (Figure 3A), we can explore the residence time at an extended force range (Figure 3B). Taking into account the known response time of the system (*t*_*DT*_ = 150) we calculate the typical residence time (*τ*) from the measured average residence time (⟨*t*⟩_*meas*_), by solving the equation 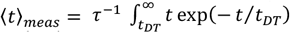. This result in ⟨*t*⟩_*meas*_ = (*t*_*DT*_ + *τ*) exp(− *t*/*t*_*DT*_), from which the approximate solution 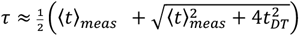 can be obtained.

### Electrophoretic Mobility Shift Assays (EMSA)

“Top” and “bottom” DNA oligonucleotides of 12, 22, and 32 bp length were purchased from IDT. The top strands were 5’-end labeled with ^32^*p* using T4 Polynucleotide Kinase (New England Biolabs). The labeled strands were then annealed, in an excess concentration, with their corresponding complementary DNA strands and purified from a gel. The DNA constructs were incubated for 3 hours on ice (∼4 ° C) with a gradient of concentrations of Egr1 in binding buffer. The samples were inserted on a native 12% polyacrylamide in 1X TBE gel while running, to ensure fast entry of the complex to the gel’s wells. Gels were run for 45 min at 200 V, ensuring that the temperature of the running buffer is maintained below 10°C. Gels were dryed (BioRad, 583) and imaged using a GE Typhoon FLA7000 phosphoimager. The fraction of bound protein was estimated using ImageLab. Curves for the fraction bound vs. Egr1 concentration were fitted to a hyperbolic binding equation to extract the dissociation constant (Figure S6F). To validate that [DNA] ≪ [Egr1], we conducted experiments at different DNA concentrations that resulted in the same binding curves. The results shown are an average of multiple gels, and the error bars are the standard error of the mean (Figure S6G).

## Author Contributions and Notes

H.K. performed the experiments and analyzed the data. S.R. prepared experimental materials and provided valuable input. H.K. and A.K. wrote the paper. P.M and A.K. supervised the research.

The authors declare no conflict of interest.

This article contains supporting information online.

## Acknowledgments

This work was supported by the Israel Science Foundation (Grants 1782/17 and 937/20 to AK, and 1850/17 to PM).

