## Supplementary Information for "Single molecule characterization of the binding kinetics of a transcription factor and its modulation by DNA sequence and methylation"

### Supplementary Discussion

**Active reannealing vs. passive binding to dsDNA**

Assuming that the DNA thermal fluctuations are in rapid equilibrium as compared to Egr1 binding, we can write predictions for the observed association rate ($k_{on}^{obs}$) for each of thes possible pathways depicted in Figure 2A:

*Pathway 1 - binding passively to the already closed DNA:*

$$k_{on}^{obs_1} = k_{on}^{1} \cdot P_{closed}$$

*Pathway 2 - binding to the open ssDNA and actively closing the DNA:*

$$k_{on}^{obs_2} = k_{on}^{2} \cdot (1 - P_{closed})$$

*Pathway 3 - binding to the open ssDNA and wait for the DNA to close spontaneously:*

$$k_{on}^{obs_3} = k_{on}^{2} \cdot k_{closed}^{2} \cdot (1 - P_{closed})$$

Here, $k_{on}^{i}$ is the intrinsic binding rate to dsDNA ($i = 1$) and ssDNA ($i = 2{,}3$), and $P_{closed}$ is the probability for the fork to be in the closed state.

**Estimation of relative isomerization and binding energies from the kinetics data**

At a fixed concentration, we can describe our system using a simple two-state model, which includes a free/loosely-bound state separated by an isomerization energetic barrier $\Delta G^{\dagger}$ from a tightly-bound one (Figure S4). $\Delta G_0$ is the difference in free energy between the states. Since, by definition, the loosely-bound state is identical for all the sites, observed differences in $k_{on}$ indicate that the sequence directly affects $\Delta G^{\dagger}$. Moreover, given that $k_{off}$ depends on $\Delta G^{\dagger} - \Delta G_0$, differences in this rate that are not correlated with $k_{on}$ indicates a different binding energy $\Delta G_0$ for each binding site. Hence, measurements of the kinetic rates allow us to calculate differences in binding and isomerization energies between two arbitrary states, 1 and 2. Since $k_{on} \sim e^{-\Delta G^{\dagger}/kT}$, the difference in isomerization energy is given by $\Delta\Delta G^{\dagger} = -kT \ln\left(k_{on}^{(1)}/k_{on}^{(2)}\right)$. Next, since $k_{off} \sim e^{-(\Delta G^{\dagger} - \Delta G_0)/kT}$, differences in binding energy can be calculated as $\Delta\Delta G_0 = kT\left[\ln\left(k_{off}^{(1)}/k_{off}^{(2)}\right) - \ln\left(k_{on}^{(1)}/k_{on}^{(2)}\right)\right]$, which, as expected, reduces to or $\Delta\Delta G_0 = kT \ln(K_D^1/K_D^2)$, where $K_D^i \equiv k_{off}^{(i)}/k_{on}^{(i)}$.

We calculate the binding energy differences for the three binding sites in the *Lhb* promoter sites, relative to the consensus, and we found that they are in the 1.5 – 3.4 kCal/mole range (Table S1). A previous work, using fluorescence-based competitive binding experiments, formulated a model for the prediction of the destabilization of Egr1 by mutations in the consensus motif, which postulates that the contributions of the individual mutations are additive(*51*). Predictions of this model for the specific sites in the *Lhb* promoter (Table S1) are not in full agreement with our experiments (Table S2). While this discrepancy may reflect the existence of interdependence between the different contributions(*81*), it could also stem from differences in the flanking sequences and the size of the constructs. In our work, we tested Egr1 binding to sites in their genomic context, or inserted in long segments that mimic them, while the previous study used a very short (12 bp) DNA construct containing almost only the binding sites(*51*). We have shown previously the important role played by the flanking sequences in modulating Egr1 binding(*39*). Some properties of DNA also depend on its length, e.g., it's bendability(*82*). Moreover, thermal melting fluctuation at the ends of a short molecule may reduce the structural integrity of the binding site. Supporting this possibility, we conducted EMSA experiments that, even when performed below 10°C, show a very strong increase in the affinity of Egr1 for the consensus site, as the size of the probed molecule is increased from 12 to 22 bp (Figure S6).

Even when inserting the sites within the same context as the consensus motif (context 601), we do not obtain the same energy changes, or even the same ratios as the prediction. Table S2 presents our calculations of $\Delta\Delta G_0$ and $\Delta\Delta G^{\dagger}$ based on our data, obtained for sites -1, -2, and -3 within different contexts, taking the consensus as our reference point. Notably, the obtained $\Delta\Delta G_0$ are different from the predictions. Although these differences seem small, the dissociation constant is exponentially related to the free energy, leading to a significant change in the expected stability of the complex. Next, the calculated $\Delta\Delta G_0$ for the three sites within different contexts also resulted in energy changes that do not agree with the prediction and they differ from each other, which implies that the effect of the flanking sequences induces a more complex sequence-binding energy relationship that postulated before(*51*)

To decouple the flanking sequence's effect, we present the energy changes for sites -1, -2, and -3, relative to the same site sequence within the *Lhb* promoter in Tables S3-5, respectively. Notably, in addition to $\Delta\Delta G_0$, we can also calculate $\Delta\Delta G^{\dagger}$, information that is inaccessible using thermodynamic assays. We see that the sequence variations also change the energy barrier height, increasing for all the tested cases, suggesting that Egr1 binds the consensus sequence with the fastest isomerization rate.

In Table S6, we present the energy changes due to the methylation of the DNA. The differences are displayed relative to the unmethylated DNA. We see a binding destabilization for the consensus site and site -2, and binding stabilization for site -3. On the other hand, we see no change in $\Delta\Delta G^{\dagger}$, suggesting that the isomerization step rate is conserved upon methylation.

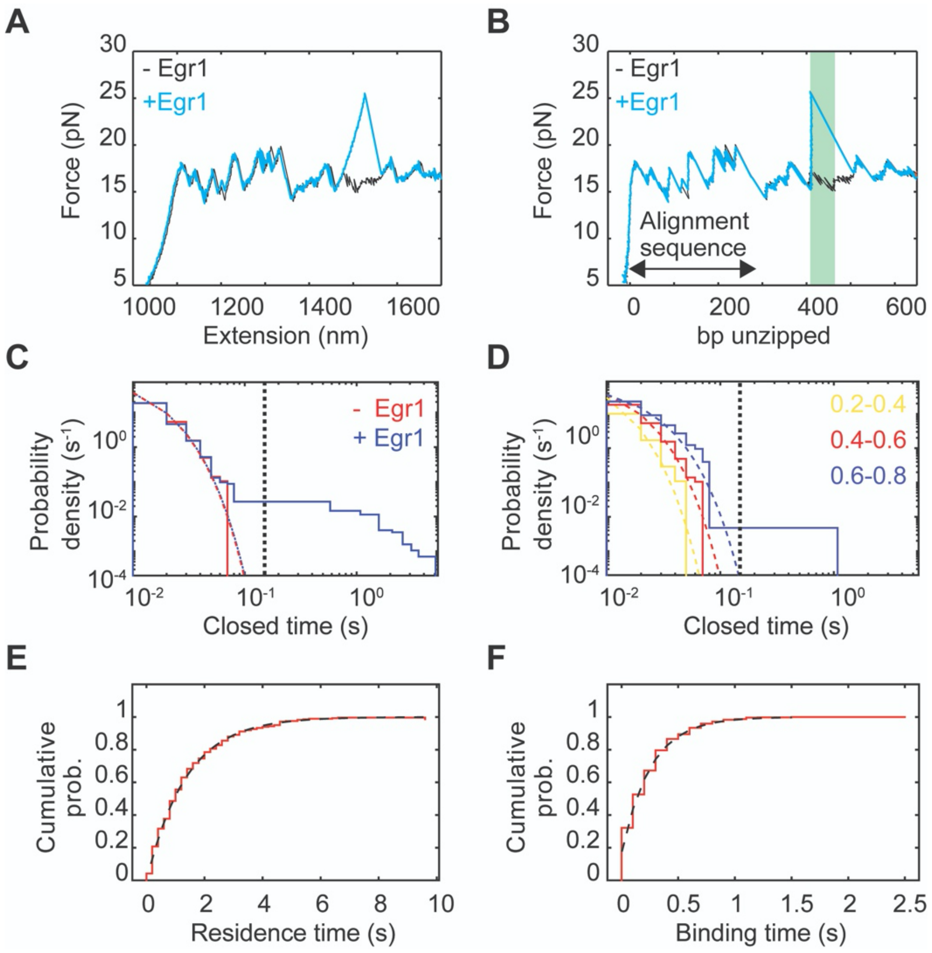


**Figure S1: (A-B)** Force vs. extension (A) and force vs. the number of bps unzipped (B), in the absence (black) and presence (blue) of Egr1. The region highlighted in green corresponds to the segment of DNA that contains the binding site. The raw data was low-pass filtered to 100 Hz (Butterworth). **(B-C)** Probability distribution function (PDF) for the closed times in the absence (red) and presence (blue) of Egr1, obtained at a $p_{closed}$ in the 0.4-0.6 range. (N = 3 and 10 experiments, 2852 and 11383 total identified times, for -Egr1 and +Egr1, respectively). Fits to a single exponential function for times shorter than 150 ms are shown as dashed lines with the same colors. The threshold time $t_{th} = 150$ ms is shown as a black dashed vertical line. **(C)** PDF of the closed times in the absence of Egr1, at different ranges of $p_{closed}$: 0.2-0.4 (yellow), 0.4-0.6 (red), and 0.6-0.8 (blue). (N=3 experiments for each range, and 2747, 2852, and 2511 identified times, respectively). **(D-E)** Binding time (D) and residence time (E) cumulative distribution functions (CDF) obtained from site -1 in the *Lhb* promoter. Exponential fits are shown as black dashed lines. (N=3 experiments, 535 and 544 identified binding and dissociation events, respectively).

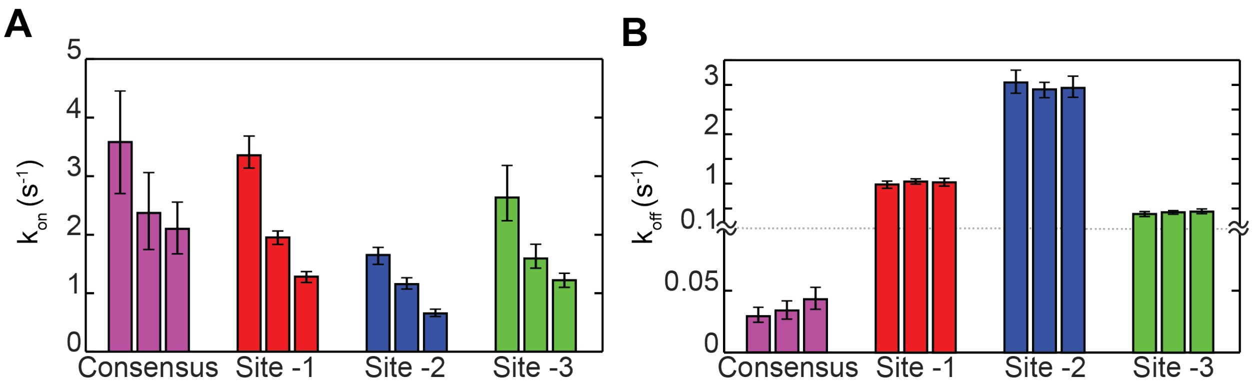


**Figure S2: (A-B)** The association (A) and dissociation (B) rates obtained for 4 sites in 3 different sessions at the same Egr1 concentration (10 nM). The rates are extracted from fitting the distributions of binding and residence time to exponential functions. The errors are shown as 95% confidence interval, obtained from bootstrapping (resampling from the data with replacement). Session 1: N=6 molecules, 114 and 93 binding and dissociation events, respectively, for the consensus; 4, 642 and 626 for site -1; 3, 344, 354 for site -2; and 3, 185, 180 for site -3. Session 2: 8, 45 and 35; 6, 1090 and 1098 for site -1; 5, 521, 543 for site -2; and 7, 368, 364 for site -3. Session 3: N=6, 100 and 83 for the consensus; 5, 511 and 508 for site -1; 4, 278, 296 for site -2; and 4, 359, 353 for site -3.

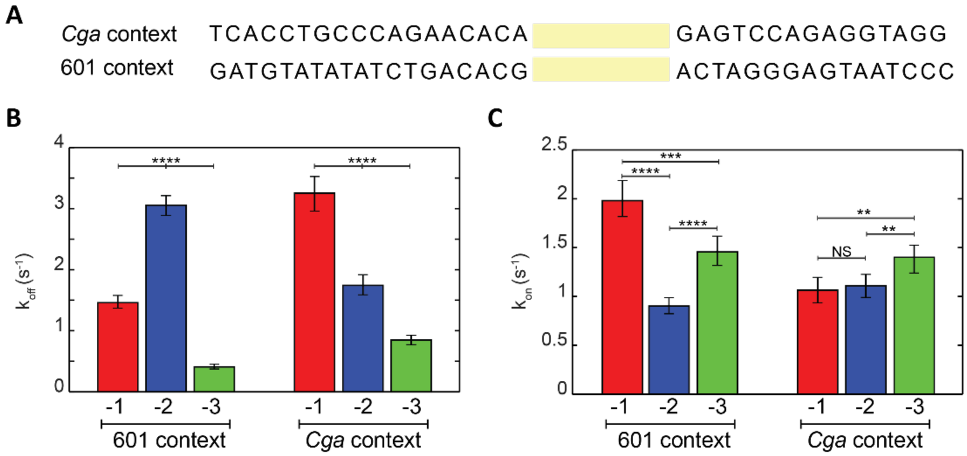


**Figure S3**: Egr1 binding kinetics for different binding sites within the same context. **(A)** The two DNA contexts that were used in the experiment. **(B-C)** The obtained dissociation (B) and association (C) rates for each of the three sites in the *Lhb* promoter, inserted into the two alternative contexts. The rates shown are extracted from fitting the binding and residence time distributions to exponential functions. The errors are shown as 95% confidence intervals, obtained by bootstrapping (resampling from the data with replacement). ****P < 0.0001, ***P < 0.001, **P < 0.01, NS P>0.05, Kolmogorov-Smirnov's t-test.

601 context: N=8 molecules, 302 and 324 binding and dissociation events, respectively, for site -1; 5, 283 and 296 for site -2; and 4, 427, 434 for site -3.

*Cga* context: 5, 497 and 513 for site -1; 5, 456 and 479 for site -2; and 6, 423, 421 for site -3.

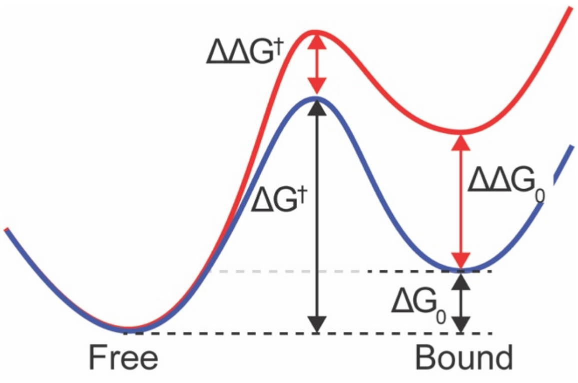


**Figure S4:** Schematic energy landscape for a two-state binding model.

| MSssI | - | - | + |
|---|---|---|---|
| FauI | - | + | + |

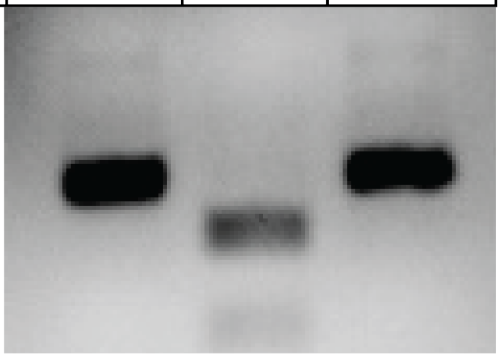

**Figure S5:** Assessment of the methylation efficiency. A DNA segment harboring a recognition site for the methylation-sensitive FauI restriction enzyme was incubated with FauI (center and right) to assess the methylation efficiency. Prior incubation with M.Sssi (rigth) inhibits the restriction.

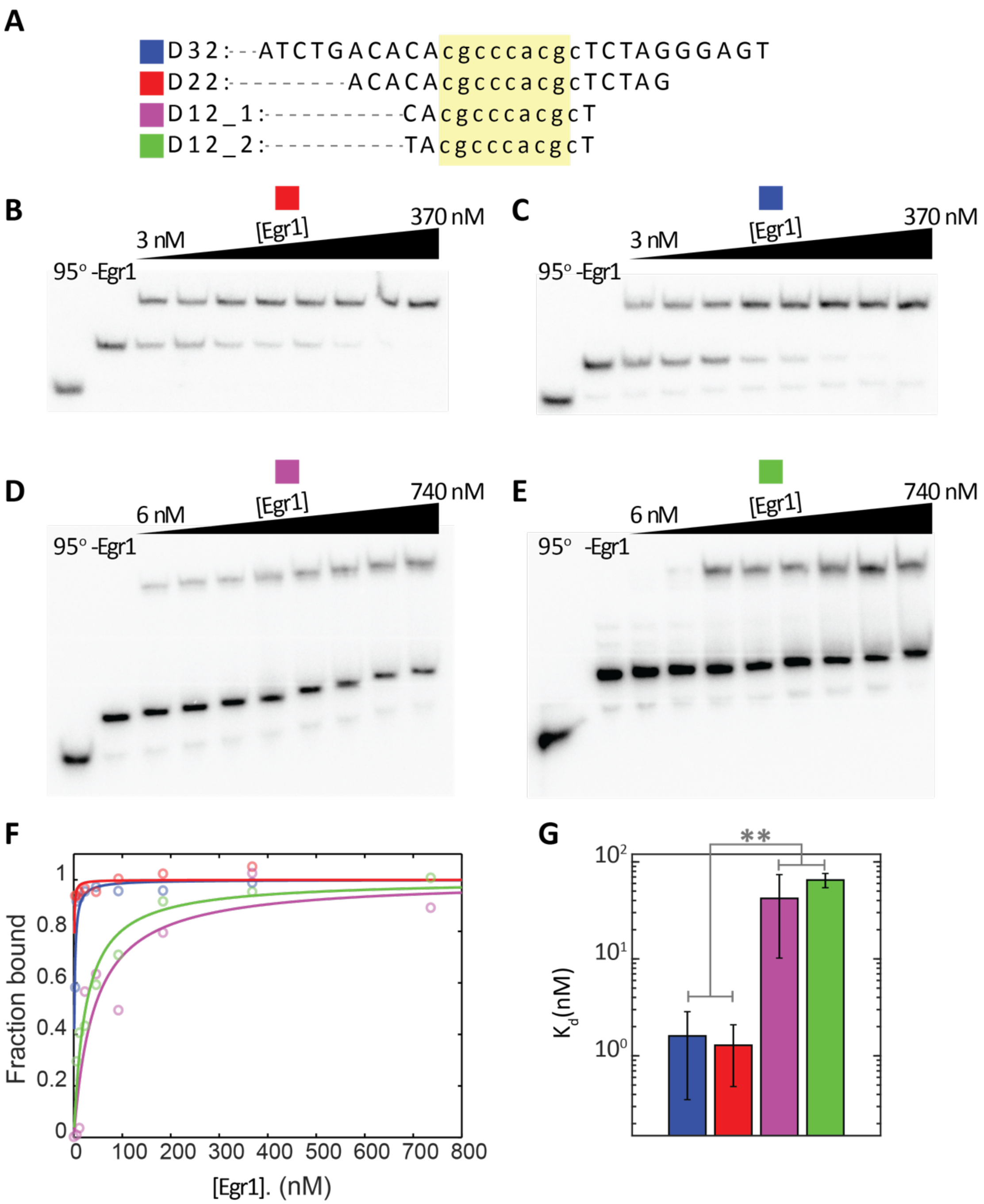


**Figure S6:** Egr1 binding affinity as a function of the DNA length. **(A)** Four DNA constructs containing the consensus binding site with different total lengths. **(B-E)** Typical EMSA gels for all the constructs. The first column shows a sample denatured by heating to 95°C, the second column a sample with no Egr1, and the following columns samples incubated at 4°C with increasing concentrations of Egr1. The gels were kept below 10°C during running. **(F)** Measured bound fraction obtained from the EMSA gels. Fits to hyperbolic binding equations are shown as solid lines. **(G)** Averaged dissociation constants, shown as the mean over multiple gels ± standard error of the mean (N=6, 6, 5, and 5 different gels for D32, D22, D12_1, and D12_2, respectively). **$p < 0.01$, two-sample Student's t-test.

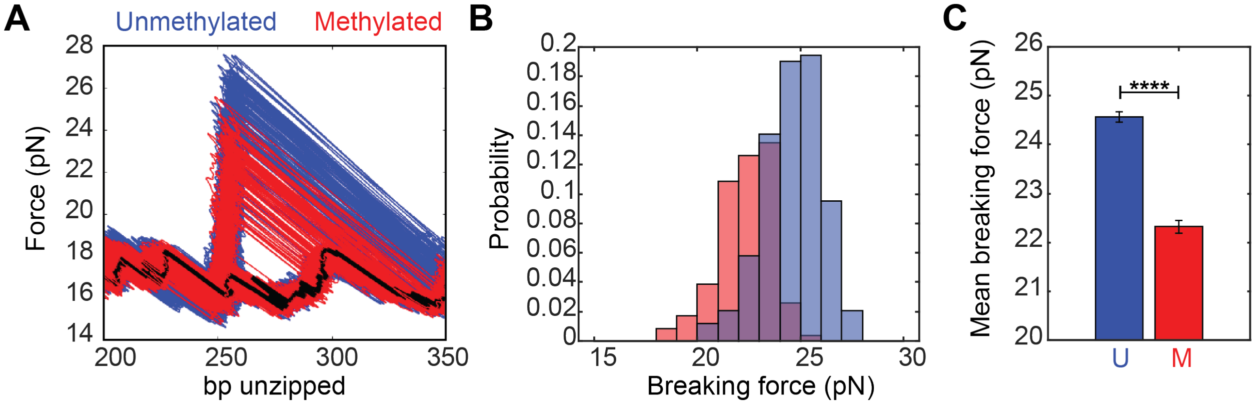


**Figure S7: (A)** Full unzipping curves of the consensus binding site in the presence of Egr1, for unmethylated (blue) and methylated (red) DNA. **(B)** Histograms of the breaking forces obtained from the unzipping experiment. **(C)** Mean breaking forces are shown as mean ± SEM. (N = 177 and 107 for unmethylated and methylated DNA, respectively). ****p < 0.0001, two-sample Student's t-test.

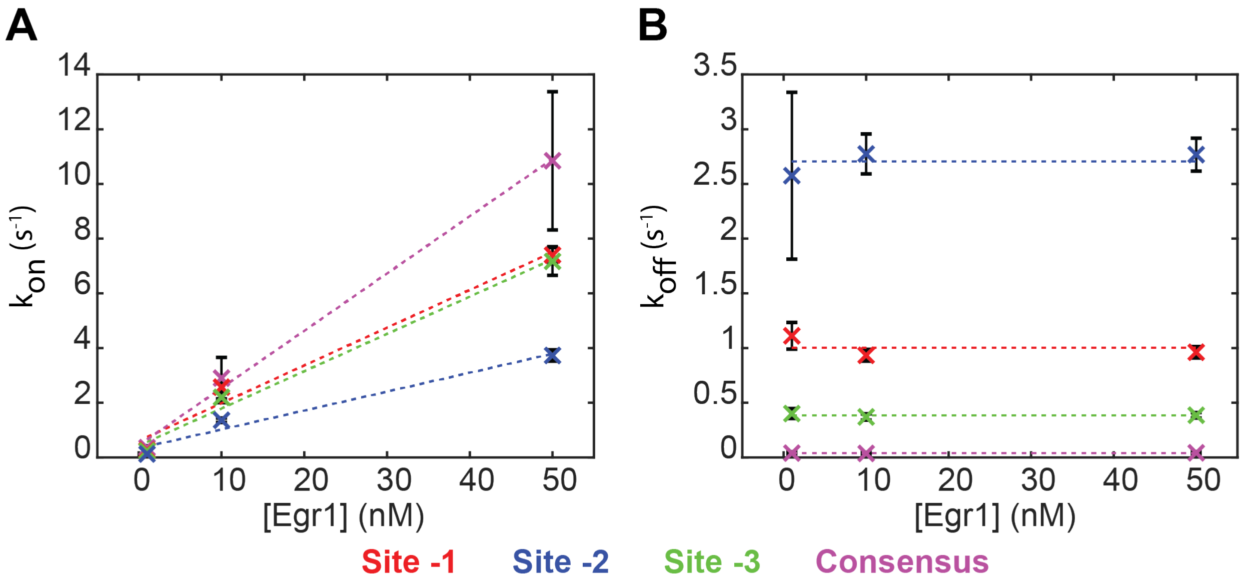


**Figure S8: (A)** Association and **(B)** dissociation rates as a function of Egr1 concentration for 4 Egr1 binding sites. Data are shown as the average of the rates extracted from the fit ± the means of the errors. (N = 3, 6, and 2 sessions for 1 nM, 10 nM, and 50 nM Egr1, respectively).

**Table S1:** Prediction of the $\Delta\Delta G_0$, relative to the consensus sequence, obtained from the model of Chattopadhyay et al. (*51*).

| | $\Delta\Delta G_0$ prediction (kcal/mol) |
|---|---|
| Site -1 | 0.6 $\pm$ 0.1 |
| Site -2 | 1.6 $\pm$ 0.1 |
| Site -3 | 0.9 $\pm$ 0.1 |

**Table S2:** Kinetic parameters obtained for various sites in various contexts and calculated $\Delta\Delta G_0$ and $\Delta\Delta G^{\dagger}$ relative to the consensus site.

| Context | Site | $k_{off}$ ($s^{-1}$) | $k_{on}$ @ 10 nM ($s^{-1}$) | $\Delta\Delta G_0$ (kcal/mol) | $\Delta\Delta G^{\dagger}$ (kcal/mol) |
|---|---|---|---|---|---|
| 601 | Consensus | 0.04 $\pm$ 0.008 | 2.1 $\pm$ 0.35 | 0 $\pm$ 0.2 | 0 $\pm$ 0.1 |
| *Lhb* | Site -1 | 1 $\pm$ 0.05 | 1.28 $\pm$ 0.08 | 2.2 $\pm$ 0.2 | -0.3 $\pm$ 0.1 |
| | Site -2 | 3 $\pm$ 0.2 | 0.65 $\pm$ 0.06 | 3.2 $\pm$ 0.2 | -0.7 $\pm$ 0.1 |
| | Site -3 | 0.43 $\pm$ 0.04 | 1.2 $\pm$ 0.1 | 1.7 $\pm$ 0.2 | -0.3 $\pm$ 0.1 |
| 601 | Site -1 | 1.5 $\pm$ 0.1 | 2 $\pm$ 0.1 | 2.1 $\pm$ 0.2 | -0.03 $\pm$ 0.1 |
| | Site -2 | 3.1 $\pm$ 0.2 | 0.9 $\pm$ 0.08 | 3 $\pm$ 0.2 | -0.5 $\pm$ 0.1 |
| | Site -3 | 0.41 $\pm$ 0.04 | 1.5 $\pm$ 0.2 | 1.6 $\pm$ 0.2 | -0.2 $\pm$ 0.1 |
| *Cga* | Site -1 | 3.7 $\pm$ 0.3 | 1.4 $\pm$ 0.1 | 2.9 $\pm$ 0.2 | -0.2 $\pm$ 0.1 |
| | Site -2 | 2.1 $\pm$ 0.2 | 1.4 $\pm$ 0.2 | 2.5 $\pm$ 0.2 | -0.2 $\pm$ 0.1 |
| | Site -3 | 1 $\pm$ 0.1 | 1.7 $\pm$ 0.2 | 2.0 $\pm$ 0.3 | -0.1 $\pm$ 0.1 |

**Table S3:** Kinetic parameters obtained for site -1 in various contexts, and calculated $\Delta\Delta G_0$ and $\Delta\Delta G^\dagger$ relative to the *Lhb* context.

| context | Site | $k_{off}$ ($s^{-1}$) | $k_{on}$ @ 10 nM ($s^{-1}$) | $\Delta\Delta G_0$ (kcal/mol) | $\Delta\Delta G^\dagger$ (kcal/mol) |
|---|---|---|---|---|---|
| *Lhb* | Site -1 | 1.03 ± 0.05 | 1.28 ± 0.08 | 0 ± 0.07 | 0 ± 0.04 |
| 601 | | 1.5 ± 0.1 | 2 ± 0.1 | 0.05 ± 0.07 | -0.3 ± 0.04 |
| *Cga* | | 3.7 ± 0.3 | 1.4 ± 0.1 | 0.7 ± 0.07 | -0.06 ± 0.04 |

**Table S4:** Kinetic parameters obtained for site -2 in various contexts, and calculated $\Delta\Delta G_0$ and $\Delta\Delta G^\dagger$ relative to the *Lhb* context.

| context | Site | $k_{off}$ ($s^{-1}$) | $k_{on}$ @ 10 nM ($s^{-1}$) | $\Delta\Delta G_0$ (kcal/mol) | $\Delta\Delta G^\dagger$ (kcal/mol) |
|---|---|---|---|---|---|
| *Lhb* | Site -2 | 3 ± 0.2 | 0.65 ± 0.06 | 0 ± 0.6 | 0 ± 0.05 |
| 601 | | 3 ± 0.2 | 0.9 ± 0.08 | -0.2 ± 0.6 | 0.18 ± 0.05 |
| *Cga* | | 2 ± 0.2 | 1.4 ± 0.2 | -0.6 ± 0.6 | 0.44 ± 0.05 |

**Table S5:** Kinetic parameters obtained for site -3 in various contexts, and calculated $\Delta\Delta G_0$ and $\Delta\Delta G^\dagger$ relative to the *Lhb* context.

| context | Site | $k_{off}$ ($s^{-1}$) | $k_{on}$ @ 10 nM ($s^{-1}$) | $\Delta\Delta G_0$ (kcal/mol) | $\Delta\Delta G^\dagger$ (kcal/mol) |
|---|---|---|---|---|---|
| *Lhb* | Site -3 | 0.43 ± 0.04 | 1.2 ± 0.1 | 0 ± 0.1 | 0 ± 0.06 |
| 601 | | 0.4 ± 0.03 | 1.5 ± 0.2 | -0.1 ± 0.1 | 0.1 ± 0.06 |
| *Cga* | | 1 ± 0.1 | 1.71 ± 0.2 | 0.3 ± 0.1 | 0.2 ± 0.06 |

**Table S6:** Kinetic parameters obtained for various sites at different methylation states, and calculated $\Delta\Delta G_0$ and $\Delta\Delta G^\dagger$ for the methylation state relative to the unmethylated one. (Highlighted area are cases with no significant change).

| Methylation status | context | Site | $k_{off}$ ($s^{-1}$) | $k_{on}$ @ 10 nM ($s^{-1}$) | $\Delta\Delta G_0$ (kcal/mol) | $\Delta\Delta G^\dagger$ (kcal/mol) |
|---|---|---|---|---|---|---|
| U | 601 | Consensus | 0.04 ± 0.01 | 0.9 ± 0.2 | 0 ± 0.3 | 0 ± 0.1 |
| M | | | 0.21 ± 0.02 | 0.9 ± 0.1 | 1 ± 0.3 | 0.02 ± 0.1 |
| U | *Lhb* | site -1 | 0.75 ± 0.05 | 3.9 ± 0.3 | 0 ± 0.09 | 0 ± 0.05 |
| M | | | 0.69 ± 0.05 | 3.8 ± 0.4 | -0.03 ± 0.09 | -0.009 ± 0.05 |
| U | | site -2 | 2.4 ± 0.2 | 2.2 ± 0.2 | 0 ± 0.09 | 0 ± 0.06 |
| M | | | 2.7 ± 0.1 | 2.5 ± 0.2 | -0.01 ± 0.09 | 0.07 ± 0.06 |
| U | | site -3 | 0.3 ± 0.04 | 3.5 ± 0.6 | 0 ± 0.2 | 0 ± 0.1 |
| M | | | 0.16 ± 0.03 | 3.8 ± 0.9 | -0.4 ± 0.2 | 0.04 ± 0.1 |

**Table S7**: Primers used to generate the binding segments.

| # | Forward/Reverse | Site | Context | Sequence |
|---|---|---|---|---|
| 1 | Forward | -1,-2,-3 | *Lhb* | ATGGCCTTGCCGGCGACACACCCC TTACTTCCAGAG |
| 2 | Reverse | -1,-2,-3 | *Lhb* | ATCACTGCGTGTCTACCCCTGACCT GGTTTTTC |
| 3 | Forward | Consensus | 601 | ATGGCCTTGCCGGCGATGTATATA TCTGACACAcgcccacgcTCTAGGGAG TAATCCCCTTG |
| 4 | Forward | -3, LM | 601 | ATGGCCTTGCCGGCGATGTATATA TCTGACACCggcccactcACTAGGGAG TAATCCCCTTG |
| 5 | Forward | -3, RM | 601 | ATGGCCTTGCCGGCAATCACCTGC CCAGAACACAggcccactcGAGTCCAG AGGTAGG |
| 6 | Forward | -3, FM | 601 | ATGGCCTTGCCGGCGATGTATATA TCTGACACGggcccactcACTAGGGAG TAATCCCCTTG |
| 8 | Forward | -1 | 601 | ATGGCCTTGCCGGCGATGTATATA TCTGACACAcacccccacTCTAGGGAG TAATCCCCTTG |
| 9 | Forward | -2 | 601 | ATGGCCTTGCCGGCGATGTATATA TCTGACACAcgcccccaaTCTAGGGAG TAATCCCCTTG |
| 10 | Forward | -3 | 601 | ATGGCCTTGCCGGCGATGTATATA TCTGACACAggcccactcTCTAGGGAG TAATCCCCTTG |
| 11 | Reverse | All | 601 | ATCACTGCGTGGTCACCGATGGAC CCTATACG |
| 12 | Forward | -1 | *Cga* | ATGGCCTTGCCGGCAATCACCTGC CCAGAACACAcacccccacGAGTCCAG AGGTAGG |
| 13 | Forward | -2 | *Cga* | ATGGCCTTGCCGGCAATCACCTGC CCAGAACACAcgcccccaaGAGTCCAG AGGTAGG |
| 12 | Forward | -3 | *Cga* | ATGGCCTTGCCGGCAATCACCTGC CCAGAACACAggcccactcGAGTCCAG AGGTAGG |
| 14 | Reverse | all | *Cga* | ATCACTGCGTGGCCATGAGCACCA TAAAGAAAATTAATGAATGC |

**Table S8**: Primers used to generate the handles.

| # | Primer name | Sequence |
|---|---|---|
| 1 | 2K BioTg F | /5'Biotin-TEG/GATCTCCAGCCAGGAACTATTGA |
| 2 | 2K bio Nt.BvCI R | GTGTCAGCTTGCCCCTCAGCGATGACCTCAGCATTT TTCGACCTGCTCTTCAGCA |
| 3 | 2K DoubleDig2 Bgl1F | ATGGCCTAGACGGCGAGCCTGGGTTTATAAGGGGA GCGGTGA |
| 4 | 2K Dig Nb.BbVCI R | TCAGCTTGCCCCTCAGCGATGACCTCAGCAAGGACC AGCGTTTTGTTGAAA |
| 5 | Oligo 5' Dig AGA | /5'Digoxigenin(NHS ester/AGGTGCCGGGACCACCCTGTAGA |
| 6 | Oligo 3' Dig/ 5' phos | /5'Phosphate/ACAGGGTGGTCCCGGCACCT/3'Digoxigeni n (NHS ester/ |
| 7 | Alignment CAC DraIII F | ATCACCACGTGGAATTCGATATCCCCGAGA |
| 8 | Alignment R DraIII | ATGCACGCAGTGACCATGGTGGTGTTTCCC |
| 9 | Hairpin oligo 5' phos | /5' Phosphate/GACTTGAGGCAATTGCCTCAAGTCTGC |

**Table S9:** DNA sequences of the DNA constructs used in the study.

| Construct | Sequence |
|---|---|
| *Lhb* | TGCCGGCGACACACCCCTTACTTCCAGAGTTCCTCCAGGCGCA<br>ATTTACTGATCAAGAAGTTTTATAGCTGAAACCACACCTATTTT<br>TGGACCCAATCCAGGCATCCTGATTAGGGGCTGGGAGACGGT<br>GGTGCACCACCTCTGGTTGGATTGAAAGCAAATTTGGAggcccact<br>cGTCAGAACCTAAGGTTGAAGCTGTGCCCTCCTATTTAGTTGTA<br>CCCAACCATCAGAGTGGGTCTGATGGACGTTTACTCCAGCAAT<br>CTGGGGGTTCAGCGAGCAGCCTGCAGTGGCCTCCCCTTTACCT<br>TGTTTCCCGTGCTTCCAATGTCAGCTAAGCCCTGACACCTGGG<br>CCGAGTGTGAGGCCAATTCACTGGGACACTGGAGCTAGTCCCT<br>GGCTTCCCTGACCTTGTCTGTGTCTcgcccccaaAGAGATTAGTGTC<br>TAGGTTACCCAAGCCTGTAGCCACTACTTAGTGGCCTTGCcaccc<br>ccacAACCCGCAGGTATAAAGCCAGGTGCCCAAGGTAGGGAAG<br>GTATCAAGAATGGAGAGGCTCCAGGTAAGATGGTAGGGCCCA<br>GGGTACTTCCCAATCCCACCAAACCCATAGATGGACAGCCTTG<br>TGACCTGGGGGTTGGAGGAGGGAGAGGAGGGGGTCTCCTAGC<br>TGGTGGACTTCAAACAGGAGATAAACAGATTTCATGGCTGGTG<br>ATGGGTCTTGAAGGACAGTTTCTGTATTATACTGAGTGGGTGC<br>AGAACCTGGGATGGAAAAACCAGGTCAGGGGTAGACACGCAG |
| 601 | ATGGCCTTGCCGGCGATGTATATATCTGACACGcainsertacACTA<br>GGGAGTAATCCCCTTGGCGGTTAAAACGCGGGGGACAGCGCG<br>TACGTGCGTTTAAGCGGTGCTAGAGCTTGCTACGACCAATTGA<br>GCGGCCTCGGCACCGGGATTCTCCAGGGCGGCCGCGTATAGG<br>GTCCATCGGTGACCACGCAGTGAT |
| *Cga* | ATGGCCTTGCCGGCAATCACCTGCCCAGAACACAcainsertacGAG<br>TCCAGAGGTAGGTAATATGATGAAATCATTTGGGGGGATTTAA<br>TTAAAGAGTAAAATATTTGTGACCACAGGAAAGAATTTTCTGC<br>ATAAATGTTGGTAGGAAAATAATTAGGCATTCATTAATTTTCT<br>TTATGGTGCTCATGGCCACGCAGTGAT |
